# Towards new cholera prophylactics and treatment: Crystal structures of bacterial enterotoxins in complex with GM1 mimics

**DOI:** 10.1101/089607

**Authors:** Julie E. Heggelund, Alasdair Mackenzie, Tobias Martinsen, Pavel Cheshev, Anna Bernardi, Ute Krengel

**Affiliations:** Department of Chemistry, University of Oslo, P.O. Box 1033, NO-0315 Blindern, Norway; Universita’ degli Studi di Milano, Dipartimento di Chimica, via Golgi 19, 20133 Milano, Italy

## Abstract

Cholera is a life-threatening disease in many countries, and new drugs are clearly needed. *C*-glycosidic antagonists may serve such a purpose. Atomic resolution crystal structures of these compounds in complexes with the cholera toxin give unprecedented atomic details of the molecular interactions, and show how the inhibitors efficiently block the GM1 binding site. These molecules are well suited for development into low-cost prophylactic drugs, due to their relatively easy syntheses and their resistance to glycolytic enzymes. One of the compounds links two toxin B-pentamers in the crystal structure, which may yield improved inhibition through the formation of toxin aggregates. These structures can spark the improved design of GM1 mimics, either alone or as multivalent inhibitors connecting multiple GM1-binding sites. Future developments may further include compounds that link the primary and secondary binding sites. Serving as decoys, receptor mimics may lessen symptoms while avoiding the use of antibiotics.

## Introduction

The secreted enterotoxins from *Vibrio cholerae* and enterotoxigenic *E. coli* (ETEC) cause millions of diarrhea episodes each year (Qadri *et al.*, 2005; Ali *et al.*, 2015). Cholera is responsible for approximately 100,000 deaths annually, a number that has been predicted to increase with climate change (Holmner *et al.*, 2010). ETEC mortality is estimated to be significantly higher, although it is difficult to determine accurate numbers due to underreporting and misdiagnosis (Qadri *et al.*, 2005). With major epidemics in recent history, there is still a requirement for rapid-acting drugs. For example, the death toll of the 2010 Haiti cholera epidemic has reached over 9000; and with the reconstruction efforts experiencing a major setback with hurricane Matthew in October 2016 (UN News Centre, 2016), the specter of cholera looms again. Except for the newly licensed live-attenuated vaccine Vaxchora (PaxVax, U.S.A.), all current vaccines require two spaced doses, and are consequently not effective in an epidemic setting. No prophylactic drugs against cholera are currently on the market.

The cholera toxin (CT) and the heat-labile enterotoxin (LT) consist of one catalytically active A-subunit bound to five non-toxic B-subunits that are arranged in a homopentamer (Merritt & Hol, 1995). The B-pentamers (CTB and LTB, respectively) are responsible for the binding to epithelial cells in the small intestine, facilitating the endocytosis of the toxin (Chinnapen *et al.*, 2007; Heggelund *et al.*, 2015). Inside the intestinal cell, the A-subunit causes a signaling cascade leading to watery diarrhea by the opening of ion channels. The resulting diarrhea can be up to 1 liter/hour, leading to life-threatening dehydration if left untreated (Sack *et al.*, 2004; Harris *et al.*, 2012). Treatment is accomplished with the application of oral rehydration therapy, but this requires medical competence and large quantities of clean water, both of which can be limited resources during an epidemic. While general antibiotics can be used against cholera infections, they are only used in serious cases and have been shown to limit the duration of the disease by 50% (Nelson *et al.*, 2011).

At present there are three cholera vaccines on the market; Shanchol (Shantha Biotechnics, India), Euvichol (EuBiologics, Korea) and Dukoral (Crucell, Netherlands), the latter of which has been shown to be effective towards both cholera and ETEC-induced diarrhea (Jelinek & Kollaritsch, 2008). All three are inactivated vaccines that have to be taken in two spaced doses, and are hence impractical in a situation where rapid protection is required, such as during a cholera outbreak. They are most frequently used for travelers from non-endemic areas, and are not effective on children under the age of 1-2 years (WHO, 2010). The only live attenuated vaccine, CVD 103-HgR (Berna Biotech, formerly Swiss Serum and Vaccine Institute, Switzerland) was taken off the market in 2003 for financial reasons (Herzog, 2016). This vaccine is currently under re-assessment, now produced in the U.S. (Vaxchora, PaxVax), and has shown to have great potential for being more effective than the other vaccines. It is also more suitable for use after outbreaks since it only requires one dose (Chen *et al.*, 2016; Harris, 2016).

The primary receptor of both CT and LT is the GM1 ganglioside (Holmgren *et al.*, 1975). The binding of CT to the GM1 oligosaccharide (GM1-os) Galβ3GalNAcβ4[NeuAcα3]Galβ4Glc (Figure 1) is one of the strongest protein-carbohydrate interactions known, with a binding constant of 43 nM (Kuziemko *et al.*, 1996; Turnbull *et al.*, 2004). Binding has been described as a “two-fingered grip”, provided by the two terminal residues, galactose (Gal) and sialic acid (NeuAc) (Merritt *et al.*, 1994). The specificity of this interaction is mainly determined by the terminal galactose residue, which is buried in a deep pocket at the rugged underside of the toxin (distant from the A-subunit). Methyl β-galactopyranoside (GalOMet) alone has been shown to bind to the CT with a *K*_D_ of 15 mM (Turnbull *et al.*, 2004). Binding of the sialic acid residue is not as strong (>200 mM), with its contribution resulting mainly from the conformational preorganization of GM1-os. CT and LT, with their five equivalent binding subunits, are known to be multivalent proteins. By binding five GM1 molecules simultaneously, the binding strength can be increased by at least an order of magnitude (Lauer *et al.*, 2002).

**Figure 1.**
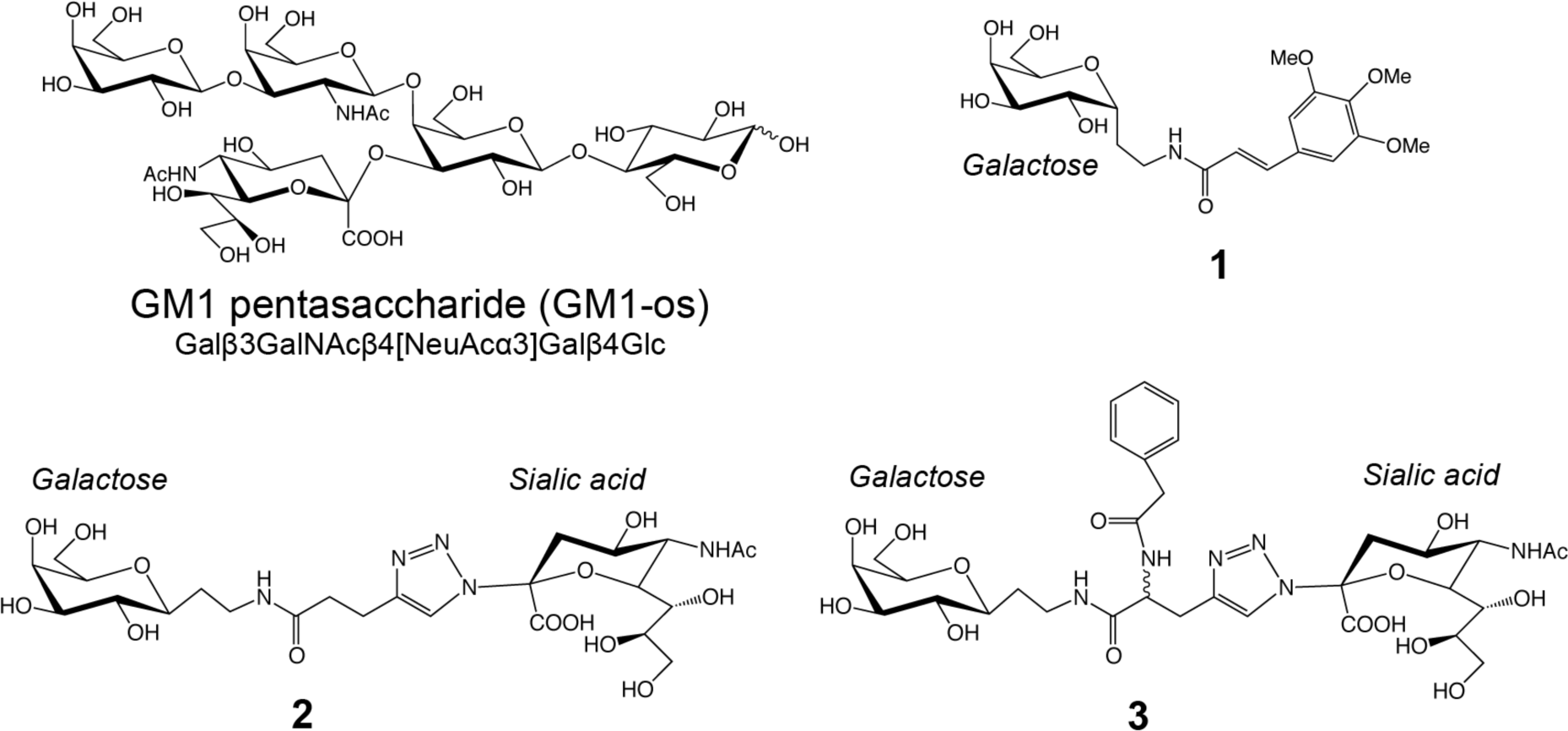
Schematic representation of GM1 pentasaccharide and inhibitors 1-3. Carbohydrate residues are labeled, and the reducing end of GM1-os is indicated by a waved line. Compound **3** was used as a mixture of two stereoisomers (*R* and *S*) at the center, indicated by a waved line.

In addition to GM1, the CT and related toxins have been shown to interact with fucosylated blood group antigens and derivatives (Holmner *et al.*, 2004; Holmner *et al.*, 2007; Heggelund *et al.*, 2012; Mandal *et al.*, 2012; Vasile *et al.*, 2014; Heggelund *et al.*, 2016). Toxin binding to the blood group H-determinant Lewis-y (Fucα2Galβ4[Fucα3]GlcNAc) characteristic of blood group O, and its blood group A-counterpart, have recently been characterized both crystallographically and by quantitative binding analyses (Heggelund *et al.*, 2016). The blood group antigens bind at the lateral side of the toxin, approximately 10 Å from the primary binding site. In all structures, the ligands are anchored to the toxin via a fucose residue. The binding strength of the Lewis-y blood group determinants is in the millimolar range, and the ligand has been suggested to serve as a secondary receptor for cell entry (Holmner *et al.*, 2004; Holmner *et al.*, 2007; Mandal *et al.*, 2012; Heggelund *et al.*, 2016). Indeed, recent cell biology experiments have shown that fucosylated carbohydrate structures can serve as functional receptors in cells in which GM1 synthesis is inhibited (Wands *et al.*, 2015). The cholera toxins come in two varieties: classical (c) and El Tor (ET). The two biotypes differ at two amino acid residues in the B-subunit, 18 and 47 (His18 and Thr47 in cCTB, and Tyr18 and Ile47 in ET CTB). While ET CTB displays reduced affinity for blood group A and B antigens (Mandal *et al.*, 2012; Heggelund *et al.*, 2016), both CT variants bind equally strongly to GM1 (Dubey *et al.*, 1990). The classical CT was by far more abundant in ancient times, and is undergoing a resurgence again today (Nair *et al.*, 2006).

The search for an effective CT inhibitor was accelerated by the publications of the crystal structures of CT and LT in the 1990’s (Sixma *et al.*, 1991; Merritt *et al.*, 1994). The major strategy in inhibitor design has been to use the terminal galactose as an anchor, for example in the promising molecule *m*-nitrophenyl-α-D-galactopyranoside (MNPG). MNPG showed enhanced binding capability compared to D-galactose, presumably through the favorable displacement of a water molecule in the binding site, enabling the direct interaction of the nitro group of MNPG to Gly33 (Merritt *et al.*, 1997). Several crystal structures were solved to investigate the potential for MNPG and its derivatives as potent inhibitors (Minke *et al.*, 2000; Pickens *et al.*, 2002; Mitchell *et al.*, 2004). Lately the search has turned towards multivalent inhibitors, using pentavalent GM1-os on different scaffolds, creating a 1:1 interaction of toxin and inhibitor (Garcia-Hartjes *et al.*, 2013; Mattarella *et al.*, 2013). An interesting approach is to “let CT fight itself” using CTB modified with GM1-os residues as penta-GM1-os-CTB neoglycoprotein inhibitors (Branson *et al.*, 2014). This potent glycoprotein inhibitor was shown to bind 1:1 to native CTB with picomolar affinity. Other potential CT inhibitors are polyphenolic compounds from grape-seed extract, suggesting that antagonists do not necessarily have to be based on carbohydrate structures (Reddy *et al.*, 2013; Cherubin *et al.*, 2016).

Despite the progress in chemo-enzymatic synthesis of GM1-os, the synthesis of this oligosaccharide is still expensive, and polyvalent GM1-os inhibitors are unlikely to result in a low-cost drug (Zuilhof, 2016). GM1 mimics have the potential to overcome this problem, and to result in materials that are relatively inexpensive and sufficiently active. Various approaches have been adopted to mimic GM1-os (Bernardi & Cheshev, 2008; Cheshev *et al.*, 2010; Ramos-Soriano *et al.*, 2013). In particular, searching for non-hydrolyzable analogs, we explored both simple *C*-galactosides (Figure 1, **1**; Podlipnik *et al.*, 2007) and bidentate ligands obtained by connecting a *C*-galactoside to a *N*-sialyltriazole residue (Figure 1, **2** and **3**; Cheshev *et al.*, 2010). These molecules are significantly simpler to synthesize than GM1-os and are stable to glycolytic enzymes, due to the absence of proper *O-*glycosidic linkages. In this paper, we describe the X-ray crystal structures of three such inhibitors (Figure 1, **1-3**), binding to the cholera toxin and the heat-labile enterotoxin. Compound **1** (Podlipnik *et al.*, 2007) is a 3,4,5-trimethoxycinnamic acid galactoconjugate that can be synthesized without the need for protective groups, and has the potential for further extensions. Compounds **2** and **3** are bidentate ligands that combine the two terminal residues of the GM1 oligosaccharide, *i.e.* galactose and sialic acid (Cheshev *et al.*, 2010).

The binding strengths of all three ligands have been measured, and found to be in the upper μM range (Podlipnik *et al.*, 2007; Cheshev *et al.*, 2010). While this is probably not enough for sufficient GM1 inhibition alone, the ligands have the potential for being combined into multivalent receptor-binding antagonists, linking several or all five binding sites.

## Results

We present four high-resolution structures of toxin-inhibitor complexes (Table 1; Figure 2 and 3). To improve the chance of successful crystallization, we used three homologous toxins: cCTB, ET CTB and porcine LTB (pLTB; with single-site mutation R13H). All three toxins have the same amino acid sequence in the primary binding site, and they also have essentially identical 3D structures. Inhibitor **1** was crystallized with ET CTB, inhibitor **2** with cCTB and ET CTB, and inhibitor **3** with pLTB R13H. The cholera toxin structures presented here were solved to atomic resolution, allowing for detailed analysis of the interactions. They are only surpassed by the recently published cCTB structure in complex with blood group determinants, solved to 1.08 Å resolution (PDB ID: 5ELB; Heggelund *et al.*, 2016). The pLTB structure is only surpassed by a structure solved to 1.3 Å (PDB ID: 1DJR; Minke *et al.*, 2000), and is the first deposited structure of the R13H variant.

**Figure 2.**
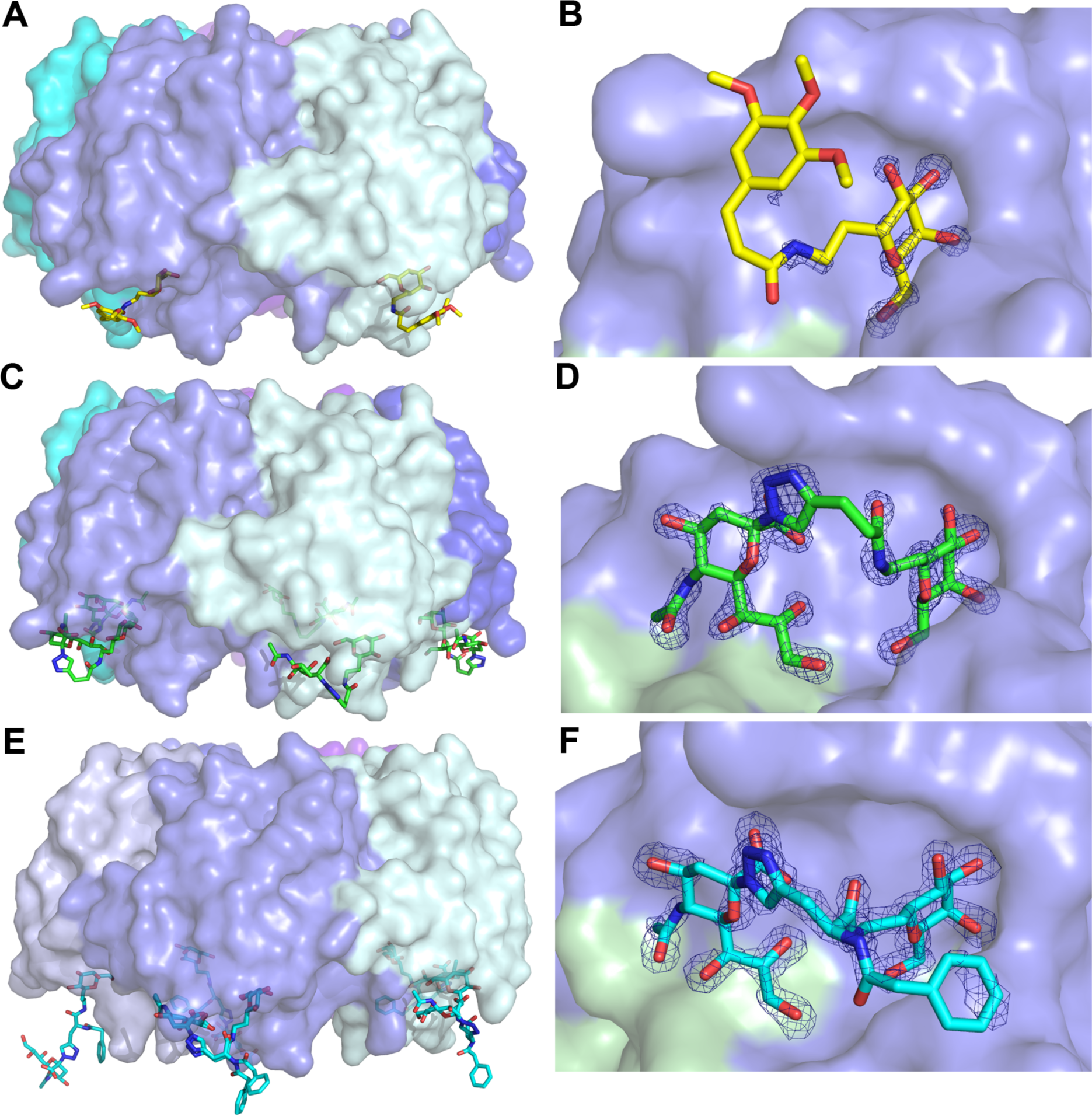
Structures of the toxin-antagonist complexes presented in this paper. The toxin B-pentamer is shown in 40% transparent surface representation, and the ligand in stick representation. Panels on the right show close-up views of the inhibitor binding sites, with σ_A_–weighted *F*_o_-*F*_c_ electron density maps contoured at 3.0 σ (generated before modeling the ligand). **(A, B)** Toxin + **1** (yellow sticks), **(C, D)** Representative structure of the toxin in complex with **2** (green sticks), **(E, F)** Toxin + **3** (cyan sticks).

**Figure 3.**
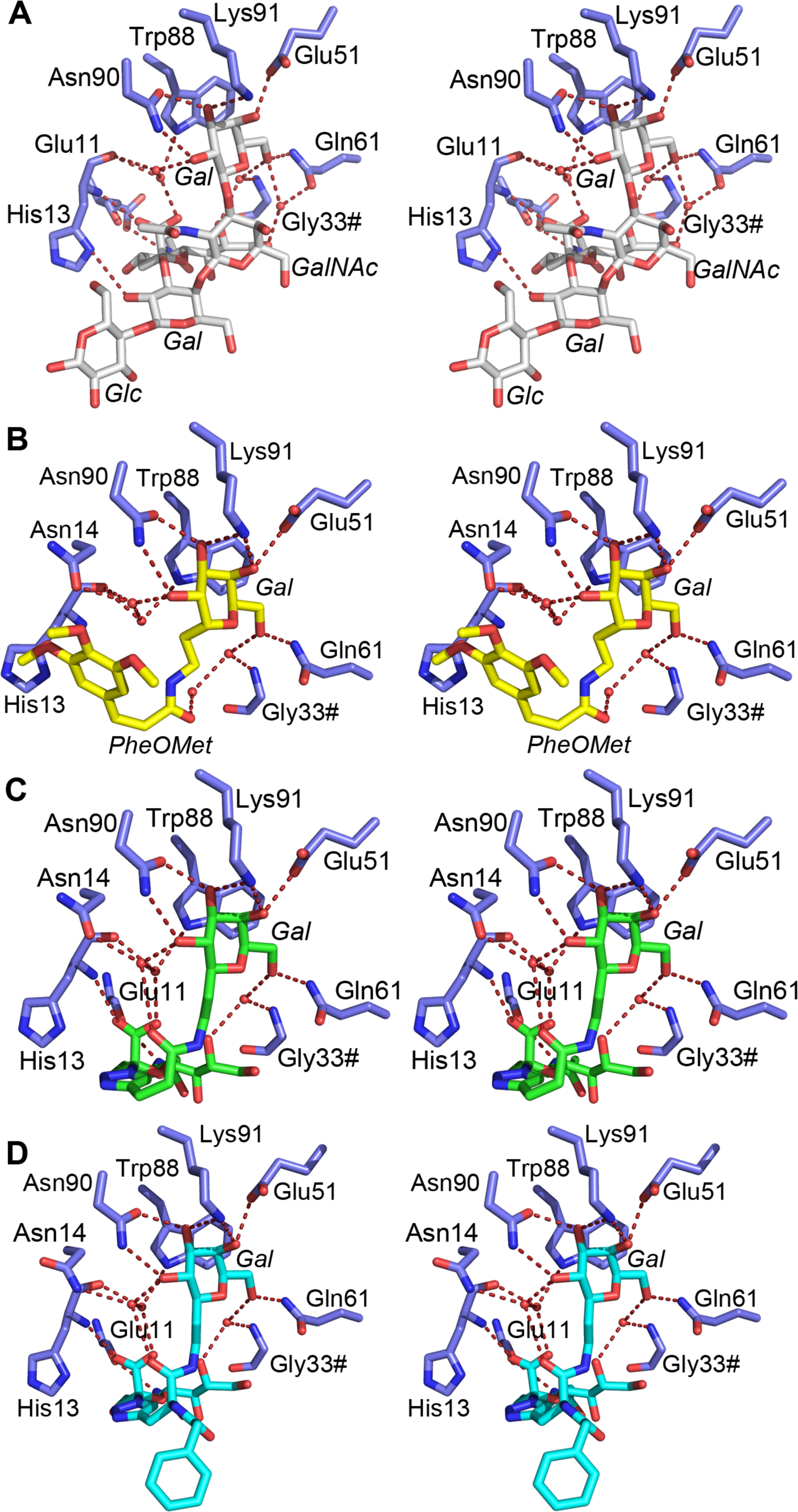
Stereoimages of the detailed carbohydrate-toxin interactions. Relevant amino acids are shown as blue sticks and labeled. Residues from neighboring subunits are indicated with a hash (#). Water molecules are shown as red spheres, and hydrogen bonds by red dashed lines (restricted to bond lengths less than 3.6 Å, and with favorable angles). Carbohydrate residues are labeled in italics. Shown are toxin complexes with **(A)** GM1-os (white sticks; PDB ID 3CHB; Merritt *et al.*, 1998), **(B) 1** (yellow sticks), **(C) 2** (green sticks), and **(D) 3** (cyan sticks). All ligands bind to the same binding site, **B-D** show structures solved in this work.

**Table 1.** Data collection and refinement statistics.

| Data set | ET CTB+ 1 | ET CTB + 2 | cCTB+ 2 | pLTB+ 3 |
| --- | --- | --- | --- | --- |
| ESRF Beam line | ID29 | ID23-1 | ID23-1 | ID14-1 |
| Space group | *C2* | *C2* | *C2* | *P2<sub>1</sub>2<sub>1</sub>2<sub>1</sub>* |
| Unit cell parameters |  |  |  |  |
| a | 102.3 | 101.9 | 101.7 | 63.2 |
| b | 66.5 | 66.3 | 66.1 | 75.9 |
| c | 82.4 | 77.4 | 76.4 | 125.2 |
| $\alpha$ | 90 | 90 | 90 | 90 |
| $\beta$ | 117.0 | 105.7 | 105.7 | 90 |
| $\gamma$ | 90 | 90 | 90 | 90 |
| Resolution (Å) | 45.6 – 1.2<br>(1.27-1.20) | 47.8 – 1.1<br>(1.19 – 1.12) | 49.0 – 1.1<br>(1.19-1.13) | 29.2-1.6<br>(1.63-1.60) |
| No. of unique reflections | 144269<br>(17960) | 139423<br>(4149) | 135800<br>(4123) | 74271<br>(2413) |
| Redundancy | 2.9<br>(2.3) | 2.7<br>(1.6) | 2.8<br>(1.7) | 2.9<br>(2.0) |
| Completeness (%) | 93.8<br>(72.4) | 74.3<br>(13.7) | 73.7<br>(13.9) | 92.9<br>(61.7) |
| $I/\sigma(I)$ | 5.55<br>(0.73) | 8.94<br>(0.70) | 6.87<br>(0.73) | 10.9<br>(3.1) |
| $R_{\text{meas}}$ | 12.1<br>(>100) | 6.5<br>(>100) | 8.4<br>(96.7) | 7.2<br>(30.6) |
| $CC_{1/2}$ | 99.4<br>(33.7) | 99.9<br>(35.9) | 99.7<br>(50.4) | 99.6<br>(90.2) |
| $R_{\text{cryst}} / R_{\text{free}}$ (%) | 19.5 / 22.8 | 17.5 / 19.8 | 19.7 / 22.9 | 14.7 / 17.6 |
| r.m.s.d bond length (Å) | 0.021 | 0.019 | 0.028 | 0.022 |
| r.m.s.d bond angles (°) | 2.15 | 1.90 | 2.50 | 2.24 |
| Average *B*-factor (Å<sup>2</sup>) |  |  |  |  |
| Backbone | 15.1 | 16.2 | 16.1 | 10.4 |
| Side chains | 18.1 | 17.7 | 17.8 | 12.9 |
| Water | 25.0 | 25.2 | 26.3 | 24.9 |
| Inhibitor | 35.6 | 19.4 | 19.2 | 15.9 |
| All atoms | 18.0 | 18.2 | 18.3 | 14.1 |
| Ramachandran plot profile (%) |  |  |  |  |
| Favored | 97.7 | 97.3 | 97.0 | 97.6 |
| Allowed | 2.3 | 2.7 | 3.0 | 2.4 |
| Disallowed | 0.0 | 0.0 | 0.0 | 0.0 |
| PDB ID | 5LZJ | 5LZG | 5LZH | 5LZI |
| Values in parentheses refer to the highest resolution shells. $R_{\text{cryst}} = \Sigma \vert \vert F_o\vert - \vert F_c\vert \vert / \Sigma \vert F_o\vert $, where $\vert F_o\vert $ and $\vert F_c\vert $ are the observed and calculated structure factor amplitudes, respectively. | | | | |

### Crystal structures of ET CTB in complex with inhibitor 1

The toxin complex with inhibitor **1** crystallized in space group *C*2, and the structure was refined to 1.2 Å (Table 1). Sufficient electron density for **1** was present in two out of five binding sites (chains C and D); two sites were occluded by crystal contacts, and the remaining site had too weak density to model the ligand with confidence. **1** binds with its galactose moiety in the galactose-binding pocket, interacting with Glu51, Gln61, Asn90 and Lys91 (Figures 2A,B and 3B). The aryl moiety seems to be disordered in the crystal structure and is not clearly bound in the hydrophobic patch, differing from previous predictions (Podlipnik & Reina, 2012). No clear electron density is visible for the ligand after the peptide bond (Figure 2B). The binding sites lacking bound ligand display less defined loop regions (residues 50-60; chains A, B and E), indicating that ligand binding causes ordering of the CTB structure.

### Crystal structures of cCTB and ET CTB in complex with inhibitor 2

Both toxin complexes with inhibitor **2** crystallized in space group *C*2, exhibiting the same crystal form as ET CTB co-crystallized with **1**. The crystal structures were solved to 1.1 Å resolution and contain one B-pentamer in the asymmetric unit. Both CT variants have well-defined electron density for inhibitor **2** in all binding sites (Figure 2D). The five crystallographically independent binding sites show unprecedented details of the inhibitor-protein interactions. As in the other structures, the galactose moiety of **2** is buried in the GM1 binding site, H-bonding with Glu51, Gln61, Asn90 and Lys91 (Figures 2C,D and 3C). The sialic acid residue has direct interactions with the backbones of Glu11 and His13. The differences to GM1-os-binding can be seen in the linker region of **2**, where the hydrogen bond of the core Gal to the side chain of His13 is no longer achievable, but rather exchanged with a water-mediated connection between the carboxyl of the inhibitor’s peptide bond to Asn14 (Figure 3A,C). The ligand binding mode was predicted by earlier modeling experiments (Cheshev *et al.*, 2010; Podlipnik & Reina, 2012).

### Crystal structure of pLTB R13H in complex with inhibitor 3

The toxin complex with inhibitor **3** was solved to a resolution of 1.6 Å in space group *P*2_1_2_1_2_1_ with one B-pentamer in the asymmetric unit. The inhibitor was present in all five binding sites (Figure 2E,F), and binds in the same manner as **2,** with the galactose residue binding to Glu51, Gln61, Asn90 and Lys91, and the sialic acid residue interacting with Glu11 and His13 (Figure 3D). The benzylamido extension, the feature distinguishing this inhibitor from **2**, is flexible and sometimes adopts different conformations. Interestingly, in two of the binding sites, the inhibitor is stretched out and links two adjacent toxin pentamers (Figure 4). In these two cases, the sialic acid moiety extends into the sialic acid binding site of another B-pentamer, creating a cross-over with another ligand. The inhibitor solution was a mixture of two stereoisomers, as indicated by the waved line in Figure 1. Both isomers displayed very similar retention factors on weak affinity chromatography with immobilized CTB (Bergström *et al.*, 2009), suggesting a similar affinity for the protein (Cheshev *et al.*, 2010). The *S*-isomer is predominantly seen in the crystal structure, such that only one of the five binding sites contains the *R*-isomer.

**Figure 4.**
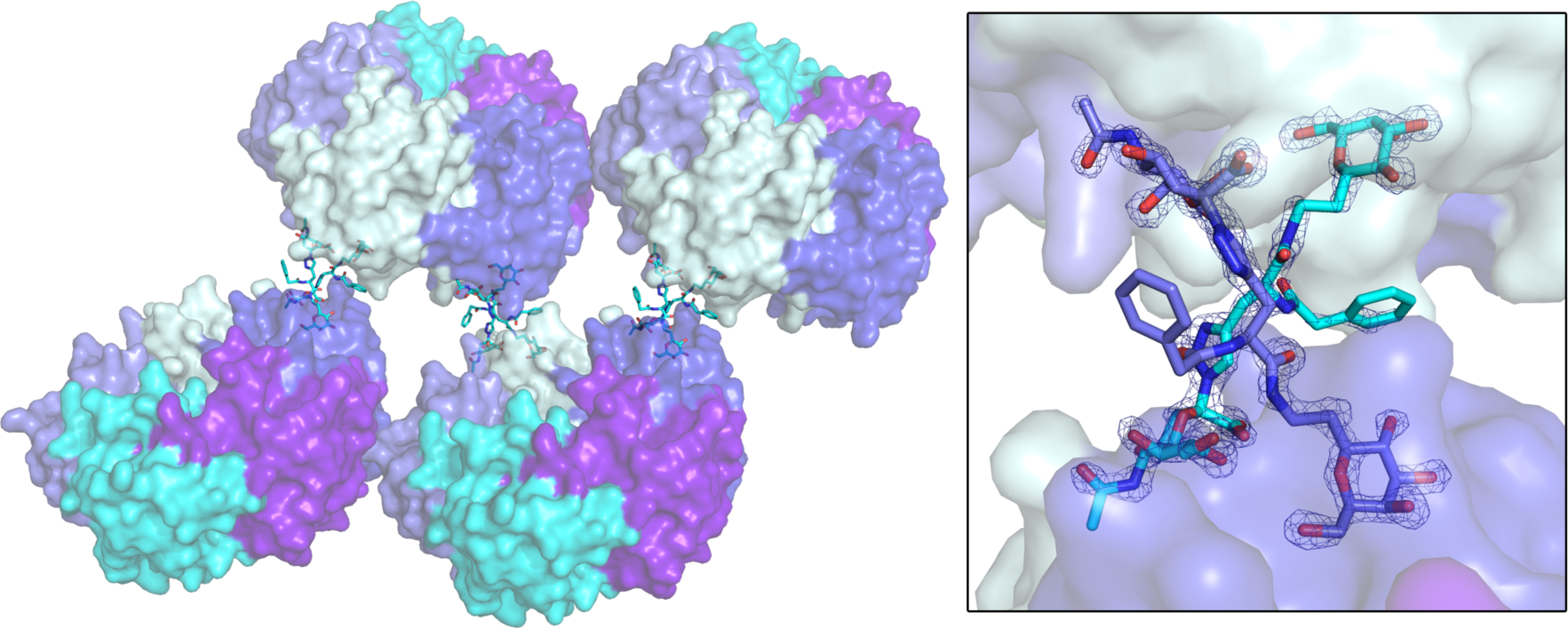
Cross-Linking of toxin B-pentamers by 3 in the crystal structure. The toxins are shown in surface representation, with the inhibitor in cyan sticks. The inset shows a close-up view of the binding interaction, with two antagonists (cyan and blue sticks) bridging two adjacent toxins. The σ_A_–weighted *F*_o_-*F*_c_ electron density map is shown for both inhibitors (contoured at 3.0 σ, generated before modeling the ligand).

## Discussion

Here we present the X-ray crystal structures of four enterotoxin inhibitor complexes at atomic resolution. As expected, all three inhibitors bind in the primary binding site, acting as decoys for the toxins’ main receptor, the GM1 ganglioside. One of the inhibitors shows the surprising capability to facilitate bridging to nearby toxins, thus potentially allowing the formation of larger aggregates. Multivalent constructs of GM1-os have been proposed before as effective antagonists of CT action in *Vibrio cholerae* infections (Zomer-van Ommen *et al.*, 2016; Zuilhof, 2016). By using these simple *C*-glycosidic mimics of GM1-os, it might be possible to simplify the organic synthesis process, which is a crucial step in the development of a low cost drug.

Ligand-induced dimerization of toxin B-pentamers has been reported previously, both for the CTB (Zhang *et al.*, 2002) and the shiga-like toxin (Kitov *et al.*, 2000). More recently, Turnbull, Zuilhof and coworkers (Sisu *et al.*, 2009) found that divalent and tetravalent analogs of GM1 were better inhibitors than pentavalent inhibitors (Zhang *et al.*, 2004; Fu *et al.*, 2015). Likewise, earlier reports found a 47,500-fold increase in binding for octavalent GM1-os dendritic glycoconjugates, resulting in an IC_50_ of 5±1×10^-11^ M (Pukin *et al.*, 2007). This is conceivably achieved through linking more than two B-pentamers together, resulting in the formation of aggregates. The bivalent inhibitor described in this paper connects receptor binding sites from different pentamers in the crystal unit cell, creating a chain of toxins (Figure 4). This would have been difficult to predict by molecular modeling, which only deals with one B-pentamer at a time. Aggregating the soluble toxin could be a very effective strategy for preventing fluid accumulation during cholera infection.

By exploiting the blood-group antigen binding site of the toxin (Figure 5), it might be possible to create even more potent inhibitors that function by promotion of the aggregation effect. It was recently shown that the two binding sites for GM1-os and blood group antigens can be occupied simultaneously (Vasile *et al.*, 2014; Heggelund *et al.*, 2016). Dual-binding site inhibitors could have the potential to induce an aggregation event by linking the primary site from one B-pentamer to the secondary site of another. Although B-pentamers in a crystal are likely positioned closer together than in the gut of a cholera-infected individual, inhibitor-induced linking of pentamers in the human gut is conceivable. The concentration of CT in human stool has been measured at 10 μg/ml (Turnbull *et al.*, 1985), and it is likely that the concentration in the small intestine is similar. Other reported inhibitors may also work by linking different binding sites in different B-pentamers, for example the high-molecular weight polysaccharide from garlic water extract that has been shown to be bioactive against CTB (Politi *et al.*, 2006).

**Figure 5.**
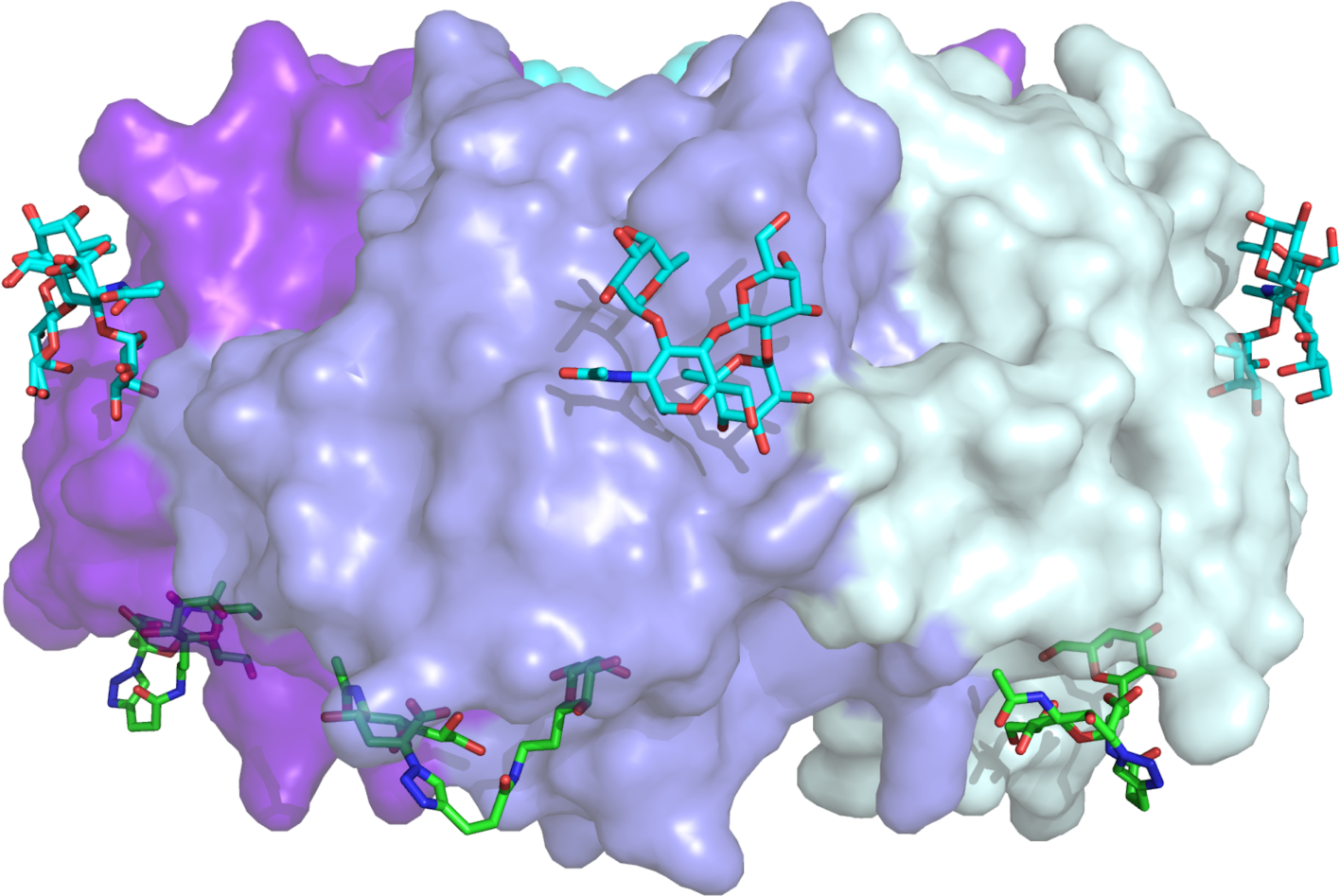
Primary and secondary binding sites of the cholera toxin. The structure of CTB with inhibitor **2**, superimposed with CTB bound to the blood group H determinant, characteristic of blood group O (PDB ID: 5ELB; Heggelund *et al.*, 2016). Only one toxin B-pentamer is shown for clarity, in 40% transparent surface representation. The blood group H determinant is shown in cyan sticks, inhibitor **2** in green sticks.

The structural insights offered in this paper can help to spark further developments in the design of potent and cost-effective cholera toxin inhibitors, especially by exploiting the promising tactic of dual-site binders. The development of such inhibitors will be crucial for making a prophylactic cure against cholera and ETEC-induced diarrhea, while at the same time avoiding antibiotics. More than 1.8 billion people use a drinking-water source contaminated with fecal matter (WHO, 2015). The development of a low-cost drug that is independent of a cold-chain delivery will have a great potential to lessen the incredible toll these diseases have on developing nations.

## Materials and Methods

### Syntheses of GM1 mimics 1-3

Inhibitor **1** (Podlipnik *et al.*, 2007) uses one component of the binding epitope, galactose, mounted on a cinnamic acid-based spacer. Inhibitor **2** (Cheshev *et al.*, 2010) mimics the GM1 oligosaccharide by connecting a sialic acid residue and galactose with a spacer consisting of a triazole group and an amide bond. Inhibitor **3** is identical to **2** except for an added benzylamido moiety.

The synthesis of **1** (compound 11) is described in (Podlipnik *et al.*, 2007), and the syntheses of **2** (compound 44) and **3** (compound 57) are described in (Cheshev *et al.*, 2010). In short, **1** was synthesized starting from easily accessible α-*C*-allyl galactoside (Bennek & Gray, 1987) that was transformed in three steps into 2-(α-D-Galactopyranosyl)ethylamine, which was finally acylated with the appropriate cinnamoyl chloride. The same starting material (α-*C*-allyl galactoside) was used for the synthesis of **2** and **3** through 2-(α-D-Galactopyranosyl)acetaldehyde. The latter was isomerized using the Massi-Dondoni protocol (Massi *et al.*, 2007) to the β-*C*-glycoside, which was transformed into 2-(β-D-Galactopyranosyl)ethylamine in two steps. Elongation with 4-pentynoic acid, followed by a click reaction with sialyl azide (Tropper *et al.*, 1992) under Sharpless conditions yielded **2**. The branched derivative **3** was obtained as **2,** by replacing pentynoic acid with commercially available racemic propargyl glycine.

### Expression of ET CTB

The gene for ET CTB (Uniprot: P01556) was previously introduced into the non-pathogenic *Vibrio* sp. 60 under an IPTG-inducible promoter (Aman *et al.*, 2001), an expression system kindly provided by Professor Timothy Hirst. The gene includes an N-terminal signal sequence directing the protein to the periplasmic space and subsequent secretion into the growth media, facilitating high expression and easy purification. The bacteria were grown in high-salt LB medium (15 g/L NaCl) supplemented with 0.1 mg/ml ampicillin at 30 °C with shaking. Expression was induced with 0.5 mM IPTG at an OD_600nm_ of 0.2, followed by protein production for 16-20 hours. The medium was separated from the bacterial pellet by centrifugation at 40,000 *g* at 20 °C, and purified further as described for all toxin B-pentamers.

### Expression of cCTB and pLTB R13H

The genes for cCTB (Uniprot: Q57193) and pLTB R13H (Uniprot: P32890 with an Arg to His substitution) were previously synthesized, cloned into the pET21b+ plasmid, and introduced into *E. coli* BL21(DE3) (Holmner *et al.*, 2011; Heggelund *et al.*, 2016). The sequence contains the signal sequence for secretion, but the protein is retained within the periplasmic space of *E. coli*, requiring purification by periplasmic lysis. The bacteria were grown in LB medium containing 0.1 mg/ml ampicillin at 37 °C until an OD_600nm_ of 0.5 was reached. After the temperature was lowered to 25ÅC, expression was induced with 0.5 mM IPTG, and protein produced for 14-18 hours. The supernatant was separated from the bacterial pellet by centrifugation at 6900 *g* and the pellet was re-suspended in ice-cold periplasmic lysis buffer (30 mM Tris pH 8, 20% (w/v) sucrose, 1 mM EDTA, 5 mM MgSO_4_), with 150 μg lysozyme per gram of cell pellet added after re-suspension. The periplasmic solution was kept cold while stirring for 10-30 minutes. The solution was centrifuged at 8500 *g* for 15 minutes and dialyzed against PBS in a Snakeskin tube (Thermo Scientific, 3500 MWCO). The resulting protein was centrifuged at 45,000 *g* for 20 minutes and purified further as described for all toxin B-pentamers.

### Purification of toxin B-pentamers

The protein was applied to a D-galactose-sepharose affinity column (Thermo Scientific) and eluted using 300 mM galactose in PBS. The fractions were concentrated using Vivaspin 20 ml concentrator tubes (5000 MWCO, PES membrane, Sartorius), and subjected to size-exclusion chromatography on a Superdex75 column mounted on an Äkta FPLC machine, pre-equilibrated with a Tris running buffer (20 mM Tris, 200 mM NaCl at pH 7.5). Fractions with toxin were dialyzed against the Tris running buffer in a Snakeskin dialysis tube (3500 MWCO, Thermo Scientific), concentrated using concentrator tubes to 3-10 mg/ml, and stored at −80 °C.

### Co-crystallization of toxins with inhibitors

Two hours before crystallization, toxins and inhibitors were mixed at a molar ration of 1:10 (B-subunit:inhibitor). Initial co-crystallization experiments were carried out at 20 °C with a crystallization robot (Oryx4, Douglas Instruments, UK). First hits were obtained in both the Morpheus screen (Gorrec, 2009), in conditions A1, A4, A9, A10 and A12, and the PGA-LM screen conditions D10 and D5 (both Molecular Dimensions). The hits were optimized using the hanging-drop vapor-diffusion technique using 24-well trays. Variations in the conditions identified by preliminary screening were subsequently explored with the use of microseeding, where seeds were prepared by crushing crystals from earlier screens with a seed bead. pLTB R13H was chosen for co-crystallization with inhibitor **3** due to its relative ease of crystallization.

The ET CTB + **1** data set was collected from a crystal found in an almost dried-out drop containing MES pH 6, 30% PEG 400, 3% PGA-LM (optimization of PGA-LM condition D5). The drop was hydrated with the same buffer, and no additional cryo-protection was necessary.

Diffraction-quality crystals were obtained for ET CTB + **2** in 0.1 M MES/Imidazole buffer, pH 6.5, with 10% PEG 1000, 10% PEG 3350, 10% MPD and 0.03 M divalent cations (optimization of the Morpheus A4 condition). cCTB + **2** crystals were obtained in 0.1 M MES/Imidazole buffer pH 6.5, with 8% PEG1000, 8% PEG 3350, 8% MPD and 0.03 M divalent cations. These crystals were cryo-protected using a buffer containing 0.1 M MES/Imidazole buffer, pH 6.5, with 12.5% PEG 1000, 12.5% PEG 3350, 12.5% MPD and 0.03 M divalent cations.

Crystals of the pLTB R13H + **3** complex were obtained from several conditions of the Clear Strategy 1 crystal screen (Molecular dimensions). Diffraction-quality crystals were grown in 0.1 M sodium cacodylate pH 6.5, 0.2 M lithium sulfate, and 15% PEG 4000, and cryo-protected using the original buffer with 25% glycerol added.

### Data collection and refinement

Samples were mounted in loops, flash-frozen in a nitrogen cryo stream, and subjected to data collection at the ESRF, Grenoble, France (Nurizzo *et al.*, 2006; Gabadinho *et al.*, 2010; de Sanctis *et al.*, 2012) (Table 1). Scaling and processing of the CTB data sets was done with XDS (Kabsch, 2010), whereas the pLTB data set was scaled and processed with Mosflm (Battye *et al.*, 2011). Diffraction cut-offs were chosen based on the assessment of CC_1/2_ (Karplus & Diederichs, 2012; Karplus & Diederichs, 2015) (Table 1). The pLTB + **3** crystal data was collected using remote access, to a resolution of 1.60 Å, which in hindsight turned out to be a very conservative cutoff with CC_1/2_ = 90.2

The structures were solved by molecular replacement using MOLREP from the CCP4 software suite (Winn *et al.*, 2011). The search model used for the CTB structures was a 1.25 ° crystal structure of cCTB (PDB ID: 3CHB; Merritt *et al.*, 1998). The pLTB R13H structure was solved using the native pLTB crystal structure (PDB ID: 1EFI; Fan *et al.*, 2001) as a search model. The search models were prepared by pruning unconserved residues and removing water molecules with the program CHAINSAW (Stein, 2008).

The inhibitors were modeled in MarvinSketch and MarvinSpace (ChemAxon.com), and the corresponding PDB‐ and library files were created using PRODRG (Schüttelkopf & van Aalten, 2004).

After initial rigid body refinement using REFMAC5 (Murshudov *et al.*, 2011), setting 5% of the reflections aside for calculating the *R*_free_, the structures were surveyed and patched with Coot (Emsley et al., 2010) and further refined. At later stages of the refinement, water molecules were manually added with Coot. The inhibitors were included last. All inhibitors were modeled with 100% occupancy after assessment of the difference electron density and *B*-factors of the ligand and the nearby protein chain.

The data sets with inhibitor **2** showed significant anisotropy, which resulted in disproportionally high *R*-factors, a common effect of anisotropy. Although the Hamilton *R* ratio test showed that the structures could also be refined with anisotropic *B*-factors, this was abandoned in favor of an isotropic *B*-factor model with TLS refinement, since anisotropic *B*-factors resulted in inferior electron density in the loop areas (residues 50-60), and made model building harder. The application of TLS refinement resulted in well-defined density also in the loop areas, along with *R*/*R*_free_ values comparable to those obtained from anisotropic *B*-factor refinement.

In the structure of cCTB in complex with **2** some of the intramolecular disulfide bridges (Cys9-Cys86) show oxidation, suggesting radiation damage. Cys9 has previously been reported to have two conformations, where one is pointing away from Cys86 and Thr15 (Merritt *et al.*, 1998; Heggelund *et al.*, 2016). However, in this structure, strong positive difference density (10 *r.m.s.d.*) was observed on the opposite site of the disulfide link, towards Thr15, indicating a modification of the sulfur atom rather than an alternative conformation of the residue. Partial oxidation of the residues was modeled by replacing the cysteines with S-oxy cysteine (CSX), with 50% occupancy of the oxygen in chains A, B and D.

The pLTB R13H + **3** data set is of high quality with close to 100% completeness, while the ET CTB + **1** data set exhibited some anisotropy. Both structures were refined with standard isotropic *B*-factors.

Figure 1 was prepared with ChemBioDraw Ultra (PerkinElmer Informatics, Inc), and figures 2-5 with PyMol (Schrödinger LLC). The coordinates and structure factors have been deposited with the Protein Data Bank (www.rcsb.org) with accession codes 5LZG, 5LZH, 5LZI and 5LZJ.

## Acknowledgments

This work was funded by the University of Oslo and the Norwegian Research Council (grant 216625), and carried out as part of the GlycoNor Consortium, University of Oslo. Funding from the University of Milano (Transition grants 2015/17) is also acknowledged. We would further like to thank Professor T. R. Hirst for providing us with the *Vibrio* sp. 60 expression system, and the staff at the ESRF for beamline support.

## References

Ali M, Nelson AR, Lopez AL, Sack DA (2015) Updated global burden of cholera in endemic countries. PLoS Negl Trop Dis 9(6):e0003832. doi: 10.1371/journal.pntd.0003832.

Aman AT, Fraser S, Merritt EA, Rodigherio C, Kenny M, Ahn M, Hol WGJ, Williams NA, Lencer WI, Hirst TR (2001) A mutant cholera toxin B subunit that binds GM1 ganglioside but lacks immunomodulatory or toxic activity. Proc Natl Acad Sci U S A 98:8536–8541. doi: 10.1073/pnas.161273098.

Battye TG, Kontogiannis L, Johnson O, Powell HR, Leslie AG (2011) iMOSFLM: a new graphical interface for diffraction-image processing with MOSFLM. Acta Crystallogr D Biol Crystallogr 67(Pt 4):271–81. doi: 10.1107/s0907444910048675.

Bennek JA, Gray GR (1987) An efficient synthesis of anhydroalditols and allylic‐ glycosides. J Org Chem 52(5):892–897. doi: 10.1021/jo00381a030.

Bergström M, Liu S, Kiick KL, Ohlson S (2009) Cholera toxin inhibitors studied with high-performance liquid affinity chromatography: a robust method to evaluate receptor-ligand interactions. Chem Biol Drug Des 73:132–141. doi: 10.1111/j.1747-0285.2008.00758.x.

Bernardi A, Cheshev P (2008) Interfering with the sugar code: Design and synthesis of oligosaccharide mimics. Chem Eur J 14(25):7434–7441. doi: 10.1002/chem.200800597.

Branson TR, McAllister TE, Garcia-Hartjes J, Fascione MA, Ross JF, Warriner SL, Wennekes T, Zuilhof H, Turnbull WB (2014) A protein-based pentavalent inhibitor of the cholera toxin B-subunit. Angew Chem Int Ed Engl 53(32):8323–7. doi: 10.1002/anie.201404397.

Chen WH, Cohen MB, Kirkpatrick BD, Brady RC, Galloway D, Gurwith M, Hall RH, Kessler RA, Lock M, Haney D, Lyon CE, Pasetti MF, Simon JK, Szabo F, Tennant S, Levine MM (2016) Single-dose live oral cholera vaccine CVD 103-HgR protects against human experimental infection with Vibrio cholerae O1 El Tor. Clin Infect Dis 62(11):1329–35. doi: 10.1093/cid/ciw145.

Cherubin P, Garcia MC, Curtis D, Britt CBT, Craft JW, Jr., Burress H, Berndt C, Reddy S, Guyette J, Zheng T, Huo Q, Quiñones B, Briggs JM, Teter K (2016) Inhibition of cholera toxin and other AB toxins by polyphenolic compounds. PLOS ONE 11(11):e0166477. doi: 10.1371/journal.pone.0166477.

Cheshev P, Morelli L, Marchesi M, Podlipnik Č, Bergström M, Bernardi A (2010) Synthesis and affinity evaluation of a small library of bidentate cholera toxin ligands: towards nonhydrolyzable ganglioside mimics. Chemistry 16(6):1951–67. doi: 10.1002/chem.200902469.

Chinnapen DJ-F, Chinnapen H, Saslowsky D, Lencer WI (2007) Rafting with cholera toxin: endocytosis and trafficking from plasma membrane to ER. FEMS Microbiol Lett 266:129–137. doi: 10.1111/j.1574-6968.2006.00545.x.

de Sanctis D, Beteva A, Caserotto H, Dobias F, Gabadinho J, Giraud T, Gobbo A, Guijarro M, Lentini M, Lavault B, Mairs T, McSweeney S, Petitdemange S, Rey-Bakaikoa V, Surr J, Theveneau P, Leonard GA, Mueller-Dieckmann C (2012) ID29: a high-intensity highly automated ESRF beamline for macromolecular crystallography experiments exploiting anomalous scattering. J Synchrotron Radiat 19(Pt 3):455–61. doi: 10.1107/s0909049512009715.

Dubey RS, Lindblad M, Holmgren J (1990) Purification of El Tor cholera enterotoxins and comparisons with classical toxin. Microbiology 136(9):1839–1847. doi: doi:10.1099/00221287-136-9-1839.

Emsley P, Lohkamp B, Scott WG, Cowtan K (2010) Features and development of Coot. Acta Crystallogr D Biol Crystallogr 66(Pt 4):486–501. doi: 10.1107/s0907444910007493.

Fan E, Merritt EA, Zhang Z, Pickens JC, Roach C, Ahn M, Hol WGJ (2001) Exploration of the GM1 receptor-binding site of heat-labile enterotoxin and cholera toxin by phenyl ring-containing galactose derivatives. Acta Crystallogr D Biol Crystallogr 57:201–212. doi: 10.1107/S0907444900016814.

Fu O, Pukin AV, van Ufford HCQ, Branson TR, Thies-Weesie DME, Turnbull WB, Visser GM, Pieters RJ (2015) Tetra‐ versus pentavalent inhibitors of cholera toxin. ChemistryOpen 4(4):471–477. doi: 10.1002/open.201500006.

Gabadinho J, Beteva A, Guijarro M, Rey-Bakaikoa V, Spruce D, Bowler MW, Brockhauser S, Flot D, Gordon EJ, Hall DR, Lavault B, McCarthy AA, McCarthy J, Mitchell E, Monaco S, Mueller-Dieckmann C, Nurizzo D, Ravelli RBG, Thibault X, Walsh MA et al. (2010) MxCuBE: a synchrotron beamline control environment customized for macromolecular crystallography experiments. J Synchrotron Radiat 17(5):700–707. doi: doi:10.1107/S0909049510020005.

Garcia-Hartjes J, Bernardi S, Weijers CA, Wennekes T, Gilbert M, Sansone F, Casnati A, Zuilhof H (2013) Picomolar inhibition of cholera toxin by a pentavalent ganglioside GM1os-calix[5]arene. Org Biomol Chem 11(26):4340–9. doi: 10.1039/c3ob40515j.

Gorrec F (2009) The MORPHEUS protein crystallization screen. J Appl Crystallogr 42(Pt 6):1035–1042. doi: 10.1107/S0021889809042022.

Harris JB, LaRocque RC, Qadri F, Ryan ET, Calderwood SB (2012) Cholera. Lancet 379(9835):2466–76. doi: 10.1016/s0140-6736(12)60436-x.

Harris JB (2016) Editorial commentary: Resurrecting a live oral cholera vaccine. Clin Infect Dis 62(11):1336–1337. doi: 10.1093/cid/ciw149.

Heggelund JE, Haugen E, Lygren B, Mackenzie A, Holmner Å, Vasile F, Reina JJ, Bernardi A, Krengel U (2012) Both El Tor and classical cholera toxin bind blood group determinants. Biochem Biophys Res Commun 418(4):731–5. doi: 10.1016/j.bbrc.2012.01.089.

Heggelund JE, Bjørnestad VA, Krengel U (2015) Vibrio cholerae and Escherichia coli heat-labile enterotoxins and beyond. In The comprehensive sourcebook of bacterial protein toxins, Alouf J, Landant D, Popoff MR (eds) pp 195–229. Elsevier.

Heggelund JE, Burschowsky D, Bjørnestad VA, Hodnik V, Anderluh G, Krengel U (2016) High-resolution crystal structures elucidate the molecular basis of cholera blood group dependence. PLoS Pathog 12(4):e1005567. doi: 10.1371/journal.ppat.1005567.

Herzog C (2016) Successful comeback of the single-dose live oral cholera vaccine CVD 103-HgR. Travel Med Infect Dis 14(4):373–377. doi: http://dx.doi.org/10.1016/j.tmaid.2016.07.003.

Holmgren J, Lönnroth I, Månsson J-E, Svennerholm L (1975) Interaction of cholera toxin and membrane GM1 ganglioside of small intestine. Proc Nat Acad Sci U S A 72(7):2520–4.

Holmner Å, Lebens M, Teneberg S, Ångström J, Ökvist M, Krengel U (2004) Novel binding site identified in a hybrid between cholera toxin and heat-labile enterotoxin: 1.9 Å crystal structure reveals the details. Structure 12:1655–1667. doi: 10.1016/j.str.2004.06.022.

Holmner Å, Askarieh G, Ökvist M, Krengel U (2007) Blood group antigen recognition by Escherichia coli heat-labile enterotoxin. J Mol Biol 371:754–764. doi: 10.1016/j.jmb.2007.05.064.

Holmner Å, Mackenzie A, Krengel U (2010) Molecular basis of cholera blood-group dependence and implications for a world characterized by climate change. FEBS Lett 584:2548–2555. doi: 10.1016/j.febslet.2010.03.050.

Holmner Å, Mackenzie A, Ökvist M, Jansson L, Lebens M, Teneberg S, Krengel U (2011) Crystal structures exploring the origins of the broader specificity of Escherichia coli heat-labile enterotoxin compared to cholera toxin. J Mol Biol 406:387–402. doi: 10.1016/j.jmb.2010.11.060.

Jelinek T, Kollaritsch H (2008) Vaccination with Dukoral against travelers’ diarrhea (ETEC) and cholera. Expert Rev Vaccines 7(5):561–7. doi: 10.1586/14760584.7.5.561.

Kabsch W (2010) XDS. Acta Crystallogr D Biol Crystallogr 66(Pt 2):125–32. doi: 10.1107/s0907444909047337.

Karplus PA, Diederichs K (2012) Linking crystallographic model and data quality. Science 336(6084):1030–1033. doi: 10.1126/science.1218231.

Karplus PA, Diederichs K (2015) Assessing and maximizing data quality in macromolecular crystallography. Curr Opin Struct Biol 34:60–68. doi: http://dx.doi.org/10.1016/j.sbi.2015.07.003.

Kitov PI, Sadowska JM, Mulvey G, Armstrong GD, Ling H, Pannu NS, Read RJ, Bundle DR (2000) Shiga-like toxins are neutralized by tailored multivalent carbohydrate ligands. Nature 403:669–672. doi: 10.1038/35001095.

Kuziemko GM, Stroh M, Stevens RC (1996) Cholera toxin binding affinity and specificity for gangliosides determined by surface plasmon resonance. Biochem (Mosc) 35(20):6375–84. doi: 10.1021/bi952314i.

Lauer S, Goldstein B, Nolan RL, Nolan JP (2002) Analysis of cholera toxin‐ganglioside interactions by flow cytometry. Biochem (Mosc) 41(6):1742–51.

Mandal PK, Branson TR, Hayes ED, Ross JF, Gavin JA, Daranas AH, Turnbull WB (2012) Towards a structural basis for the relationship between blood group and the severity of El Tor cholera. Angew Chem Int Ed Engl 51(21):5143–6. doi: 10.1002/anie.201109068.

Massi A, Nuzzi A, Dondoni A (2007) Microwave-assisted organocatalytic anomerization of α-C-glycosylmethyl aldehydes and ketones. J Org Chem 72(26):10279–10282. doi: 10.1021/jo701959b.

Mattarella M, Garcia-Hartjes J, Wennekes T, Zuilhof H, Siegel JS (2013) Nanomolar cholera toxininhibitors based on symmetrical pentavalent ganglioside GM1os-sym‐ corannulenes. Org Biomol Chem 11(26):4333–4339. doi: 10.1039/C3OB40438B.

Merritt EA, Sarfaty S, van den Akker F, L’Hoir C, Martial JA, Hol WGJ (1994) Crystal structure of cholera toxin B-pentamer bound to receptor GM1 pentasaccharide. Protein Sci 3:166–175. doi: 10.1002/pro.5560030202.

Merritt EA, Hol WGJ (1995) AB5 toxins. Curr Opin Struct Biol 5:165–171.

Merritt EA, Sarfaty S, Feil IK, Hol WGJ (1997) Structural foundation for the design of receptor antagonists targeting Escherichia coli heat-labile enterotoxin. Structure 5:1485–1499.

Merritt EA, Kuhn P, Sarfaty S, Erbe JL, Holmes RK, Hol WGJ (1998) The 1.25 Å resolution refinement of the cholera toxin B-pentamer: evidence of peptide backbone strain at the receptor-binding site. J Mol Biol 282:1043–1059. doi: 10.1006/jmbi.1998.2076.

Minke WE, Pickens JC, Merritt EA, Fan E, Verlinde CLMJ, Hol WGJ (2000) Structure of m-carboxyphenyl-α-D-galactopyranoside complexed to heat-labile enterotoxin at 1.3 Å resolution: surprising variations in ligand-binding modes. Acta crystallographica D 56:795–804. doi: 10.1107/S090744490000514X.

Mitchell DD, Pickens JC, Korotkov KV, Fan E, Hol WGJ (2004) 3,5-substituted phenyl galactosides as leads in designing effective cholera toxin antagonists; synthesis and crystallographic studies. Bioorg Med Chem 12:907–920. doi: 10.1016/j.bmc.2003.12.019.

Murshudov GN, Skubak P, Lebedev AA, Pannu NS, Steiner RA, Nicholls RA, Winn MD, Long F, Vagin AA (2011) REFMAC5 for the refinement of macromolecular crystal structures. Acta Crystallogr D Biol Crystallogr 67(Pt 4):355–67. doi: 10.1107/s0907444911001314.

Nair GB, Qadri F, Holmgren J, Svennerholm A-M, Safa A, Bhuiyan NA, Ahmad QS, Faruque SM, Faruque ASG, Takeda Y, Sack DA (2006) Cholera due to altered El Tor strains of Vibrio cholerae O1 in Bangladesh. J Clin Microbiol 44:4211–4213. doi: 10.1128/JCM.01304-06.

Nelson EJ, Nelson DS, Salam MA, Sack DA (2011) Antibiotics for both moderate and severe cholera. N Engl J Med 364(1):5–7. doi: doi:10.1056/NEJMp1013771.

Nurizzo D, Mairs T, Guijarro M, Rey V, Meyer J, Fajardo P, Chavanne J, Biasci J-C, McSweeney S, Mitchell E (2006) The ID23-1 structural biology beamline at the ESRF. J Synchrotron Radiat 13(3):227–238. doi: doi:10.1107/S0909049506004341.

Pickens JC, Merritt EA, Ahn M, Verlinde CLMJ, Hol WGJ, Fan E (2002) Anchor‐based design of improved cholera toxin and E. coli heat-labile enterotoxin receptor binding antagonists that display multiple binding modes. Chem Biol 9:215–224.

Podlipnik Č, Velter I, La Ferla B, Marcou G, Belvisi L, Nicotra F, Bernardi A (2007) First round of a focused library of cholera toxin inhibitors. Carbohydr Res 342(12-13):1651–60. doi: 10.1016/j.carres.2007.06.006.

Podlipnik Č, Reina JJ (2012) Structure based design of cholera toxin antagonists. In Cholera, Gowder S (ed) pp 177–200. inTech. doi: 10.5772/37635.

Politi M, Alvaro-Blanco J, Groves P, Prieto A, Leal JA, Cañada FJ, Jiménez-Barbero J (2006) Screening of garlic water extract for binding activity with cholera toxin B pentamer by NMR spectroscopy – an old remedy giving a new surprise. Eur J Org Chem 2006(9):2067–2073. doi: 10.1002/ejoc.200500875.

Pukin AV, Branderhorst HM, Sisu C, Weijers CAGM, Gilbert M, Liskamp RMJ, Visser GM, Zuilhof H, Pieters RJ (2007) Strong Inhibition of cholera toxin by multivalent GM1 derivatives. ChemBioChem 8(13):1500–1503. doi: 10.1002/cbic.200700266.

Qadri F, Svennerholm AM, Faruque AS, Sack RB (2005) Enterotoxigenic Escherichia coli in developing countries: epidemiology, microbiology, clinical features, treatment, and prevention. Clin Microbiol Rev 18(3):465–83. doi: 10.1128/cmr.18.3.465-483.2005.

Ramos-Soriano J, Niss U, Angulo J, Angulo M, Moreno-Vargas AJ, Carmona AT, Ohlson S, Robina I (2013) Synthesis, biological evaluation, WAC and NMR studies of S-galactosides and non-carbohydrate ligands of cholera toxin based on polyhydroxyalkylfuroate moieties. Chem Eur J 19(52):17989–18003. doi: 10.1002/chem.201302786.

Reddy S, Taylor M, Zhao M, Cherubin P, Geden S, Ray S, Francis D, Teter K (2013) Grape extracts inhibit multiple events in the cell biology of cholera intoxication. PLOS ONE 8(9):e73390. doi: 10.1371/journal.pone.0073390.

Sack DA, Sack RB, Nair GB, Siddique AK (2004) Cholera. Lancet 363:223–233. doi: 10.1016/S0140-6736(03)15328-7.

Schüttelkopf AW, van Aalten DM (2004) PRODRG: a tool for high-throughput crystallography of protein-ligand complexes. Acta Crystallogr D Biol Crystallogr 60(Pt 8):1355–63. doi: 10.1107/s0907444904011679.

Sisu C, Baron AJ, Branderhorst HM, Connell SD, Weijers CAGM, de Vries R, Hayes ED, Pukin AV, Gilbert M, Pieters RJ, Zuilhof H, Visser GM, Turnbull WB (2009) The influence of ligand valency on aggregation mechanisms for inhibiting bacterial toxins. ChemBioChem 10(2):329–337. doi: 10.1002/cbic.200800550.

Sixma TK, Pronk SE, Kalk KH, Wartna ES, van Zanten BAM, Witholt B, Hol WGJ (1991) Crystal structure of a cholera toxin-related heat-labile enterotoxin from E. coli. Nature 351:371–377.

Stein N (2008) CHAINSAW: a program for mutating pdb files used as templates in molecular replacement. J Appl Crystallogr 41(3):641–643. doi: doi:10.1107/S0021889808006985.

Tropper FD, Andersson FO, Braun S, Roy R (1992) Phase transfer catalysis as a general and stereoselective entry into glycosyl azides from glycosyl halides. Synthesis 1992(07):618–620. doi: 10.1055/s-1992-26175.

Turnbull PC, Lee JV, Miliotis MD, Still CS, Isaäcson M, Ahmad QS (1985) In vitro and in vivo cholera toxin production by classical and El Tor isolates of Vibrio cholerae. J Clin Microbiol 21(6):884–90.

Turnbull WB, Precious BL, Homans SW (2004) Dissecting the cholera toxin-ganglioside GM1 interaction by isothermal titration calorimetry. J Am Chem Soc 126(4):1047–54. doi: 10.1021/ja0378207.

UN News Centre. Haiti: UN emergency fund allocates $5 million to kick-start assistance in wake of Hurricane Matthew 2016 [20. October 2016]. www.un.org/apps/news/story.asp?NewsID=55245.

Vasile F, Reina JJ, Potenza D, Heggelund JE, Mackenzie A, Krengel U, Bernardi A (2014) Comprehensive analysis of blood group antigen binding to classical and El Tor cholera toxin B-pentamers by NMR. Glycobiology 24(8):766–78. doi: 10.1093/glycob/cwu040.

Wands AM, Fujita A, McCombs JE, Cervin J, Dedic B, Rodriguez AC, Nischan N, Bond MR, Mettlen M, Trudgian DC, Lemoff A, Quiding-Jarbrink M, Gustavsson B, Steentoft C, Clausen H, Mirzaei H, Teneberg S, Yrlid U, Kohler JJ (2015) Fucosylation and protein glycosylation create functional receptors for cholera toxin. Elife 4. doi: 10.7554/eLife.09545.

WHO. Weekly Epidemiological Report 2010. http://www.who.int/wer/2010/wer8513.pdf?ua=1.

WHO. Cholera fact sheet 391 2015. http://www.who.int/mediacentre/factsheets/fs391/en/.

Winn MD, Ballard CC, Cowtan KD, Dodson EJ, Emsley P, Evans PR, Keegan RM, Krissinel EB, Leslie AG, McCoy A, McNicholas SJ, Murshudov GN, Pannu NS, Potterton EA, Powell HR, Read RJ, Vagin A, Wilson KS (2011) Overview of the CCP 4 suite and current developments. Acta Crystallogr D Biol Crystallogr 67(Pt 4):235–42. doi: 10.1107/s0907444910045749.

Zhang Z, Merritt EA, Ahn M, Roach C, Hou Z, Verlinde CLMJ, Hol WGJ, Fan E (2002) Solution and crystallographic studies of branched multivalent ligands that inhibit the receptor-binding of cholera toxin. J Am Chem Soc 124:12991–12998.

Zhang Z, Pickens JC, Hol WGJ, Fan E (2004) Solution‐ and solid-phase syntheses of guanidine-bridged, water-soluble linkers for multivalent ligand design. Org Lett 6(9):1377–1380. doi: 10.1021/ol049835v.

Zomer-van Ommen DD, Pukin AV, Fu O, Quarles van Ufford LHC, Janssens HM, Beekman JM, Pieters RJ (2016) Functional characterization of cholera toxin inhibitors using human intestinal organoids. J Med Chem 59(14):6968–6972. doi: 10.1021/acs.jmedchem.6b00770.

Zuilhof H (2016) Fighting cholera one-on-one: the development and efficacy of multivalent cholera-toxin-binding molecules. Acc Chem Res 49(2):274–285. doi: 10.1021/acs.accounts.5b00480.

